# Measuring Biotherapeutic Viscosity and Degradation On-Chip with Particle Diffusometry

**DOI:** 10.1101/179101

**Authors:** K.N. Clayton, D. Lee, S.T. Wereley, T. L. Kinzer-Ursem

## Abstract

In absence of efficient ways to test drug stability and efficacy, pharmaceuticals that have been stored outside of set temperature conditions are destroyed, often at great cost. This is especially problematic for biotherapeutics, which are highly sensitive to temperature fluctuations. Current platforms for assessing the stability of protein-based biotherapeutics in high throughput and in low volumes are unavailable outside of research and development laboratories and are not efficient for use in production, quality control, distribution, or clinical settings. In these alternative environments, microanalysis platforms could provide significant advantages for the characterization of biotherapeutic degradation. Here we present particle diffusometry (PD), a new technique to study degradation of biotherapeutic solutions. PD uses a simple microfluidic chip and microscope setup to calculate the Brownian motion of particles in a quiescent solution using a variation of particle image velocimetry (PIV) fundamentals. We show that PD can be used to measure the viscosity of protein solutions to discriminate intact protein from degraded samples as well as to determine the change in viscosity as a function of therapeutic concentration. PD viscosity analysis is applied to two particularly important biotherapeutic preparations: insulin, a commonly used protein for diabetic patients, and monoclonal antibodies which are an emerging class of biotherapeutics used to treat a variety of diseases such as autoimmune disorders and cancer. PD-based characterization of solution viscosity is a new tool for biotherapeutic analysis, and owing to its easy setup could readily be implemented at key points of the pharmaceutical delivery chain and in clinical settings.

## Introduction

Biotherapeutic development is a multi-billion-dollar industry in response to the growing need for new disease treatments.^1^ Analyzing the stability and efficacy of protein-based biotherapeutics presents unique challenges for the pharmaceutical industry due to the propensity of these biomolecules to denature from environmental factors, such as storage conditions.^2^ For example, for most of the 422 million people worldwide who suffer from type I or type II diabetes, insulin is a critical biotherapeutic used to maintain consistent glucose levels within the blood.^3–5^ Monitoring insulin efficacy is critical considering that its activity diminishes due to both temperature and time.^6^ Indeed, maintaining the native protein folding of insulin is essential for biotherapeutic function.^7^ Despite the large global insulin market size (expected to reach $53.04 billion by 2022^8^), there is currently no method in which distributors, clinics, and patients can determine whether an insulin prescription is degraded. Present measures to determine the effectiveness of insulin from the patient perspective include (1) patients returning their prescription to the manufacturer for examination, (2) trial injection of the drug and examining the outcome, or (3) simply purchasing a new prescription. These current approaches are either inadequate, unsafe, or incur unnecessary financial burden. Additionally, drug manufacturers are required to destroy biotherapeutic products when the cold chain distribution is disrupted, which in turn elevates manufacturing costs.

Monoclonal antibodies (mAbs) also make up a large percent of the biotherapeutic market. mAbs are used as therapeutic treatments for cancer and autoimmune disorders and the market will grow to $125 billion by 2020.^9^ mAbs are provided to patients in small doses (1 mL) at high concentrations (100-150 mg/mL). Producing the mAbs at such concentrated dosages results in highly viscous solutions, which lowers manufacturability of the drug and requires larger gauge needles for biotherapeutic injection. To alleviate these downstream issues, biopharmaceutical companies are often screening for mAb formulations with lower viscosity during the research and development process. Newly produced mAbs are often made in low sample volumes and are therefore not compatible with many common rheology techniques, such as cone and plate and capillary rheometers; though recently researchers have started to apply multiple particle tracking (MPT) techniques to measure mAb viscosity.^10,11^ Measurement systems for investigating protein folding state and biotherapeutic viscosity are not currently automated in a way in that is translatable to many points in the biotherapeutic delivery chain. Engineering a simple micro-scale device to study protein degradation and viscosity could provide distributors, and even clinicians with a monitoring device that could be used to track the efficacy of biotherapeutics.

Current methods used to analyze the protein-folding state of a biotherapeutic are technologically non-translatable from research and development laboratories to distribution points without extensive innovation (Fig. 1). Native PAGE is a gel electrophoresis method that is widely used to provide information on protein electrophoretic mobility, folding state, and sample purity.^12^ It requires specialized equipment and technical training to run and analyze the results and can take at least 30 minutes to perform, although cutting edge electrophoretic techniques in microfluidic formats are being developed.^13,14^ Circular dichroism (CD) is a spectral method using a polarized light source to study characteristic signatures of protein conformation.^15^ This technique has been integral in furthering the investigation of protein structure and molecular interactions and requires advanced optics in its set-up. Similar to native PAGE, CD involves specialized equipment to run the assay and user training to interpret the results, leaving the technique to be predominantly used by structural biologists rather than quality control personnel, patients or clinicians. Other methods used to assess the ability of a therapeutic to induce a cellular response (defined as the activity or efficacy) involve precise reagent handling and luminescence or absorbance readouts that require expert training.^16^

**Fig. 1.**
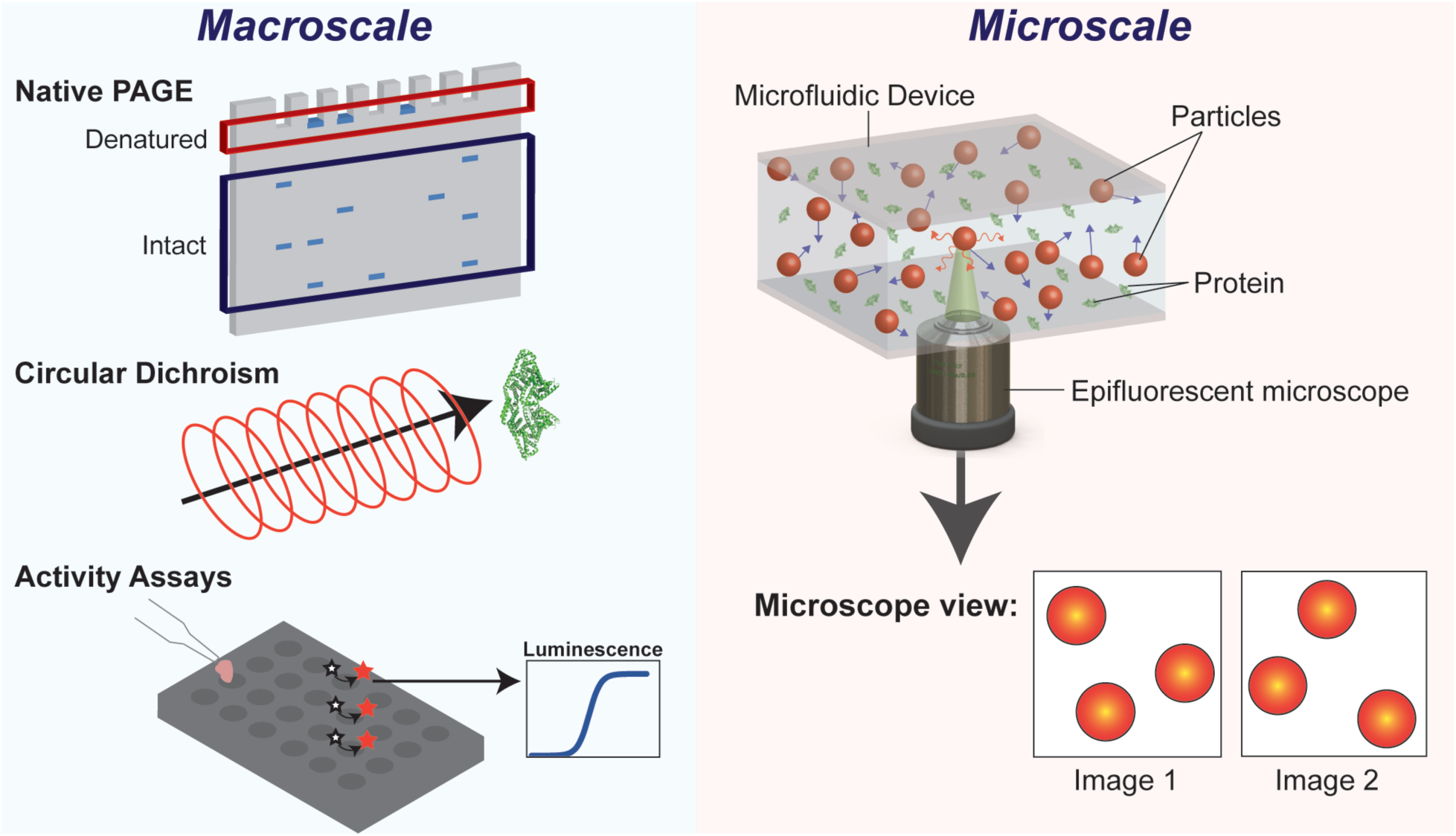
Macroscale and microscale investigations of protein structure. Gold standard macroscale systems for measuring protein folding state involve methods such as native PAGE, circular dichroism, and activity assays. Conversely, PD, a microscale system, involves imaging particles suspended in a protein solution and correlating the motion to determine sample viscosity, and therefore protein folding state.

We have recently developed the technique called Particle Diffusometry that images the Brownian motion of particles in a quiescent fluid and calculates the average diffusion coefficient of those particles. We have previously shown that PD is a statistically robust technique that is sensitive enough to detect small changes in nanoparticle size.^17^ PD correlates particle displacement across 100 sequentially-acquired images to statistically determine a diffusion coefficient in a short amount of time (typically < 8 seconds.) Since the technique fundamentally measures a particle diffusion coefficient, we can use the Stokes-Einstein equation to calculate changes in solution temperature,^18,19^ particle size,^17^ or solution viscosity. In this work, we use PD to examine how intact and denatured proteins each differently influence the viscosity of solutions by measuring intact and denatured bovine serum albumin (BSA) and insulin proteins. Further, biotherapeutic solutions typically comprise of some fraction of efficacious proteins (intact) and some fraction of denatured proteins due to accidental exposure to non-optimal conditions during distribution from the manufacturer to the patient, aging of the prescription, or other external factors.^20,21^ We study this effect by combining intact and degraded insulin in controlled ratios and study the resulting effect on solution viscosity. Finally, we investigate the effect of varying concentrations of intact mAb on solution viscosity for downstream applications in pharmaceutical research and development.

### Theory

In particle diffusometry, the diffusion coefficient of particles in solution is calculated using correlation analysis.^17^ First, a series of images of particles undergoing Brownian motion in a quiescent solution is captured using a simple microscopy setup (Fig. 1). Each individual image is then partitioned into many smaller interrogation windows. The area of the interrogation windows is defined such that 8-10 particles are situated within each interrogation window. To perform cross-correlation on the interrogation window, we correlate a first image, at time *t*, with a second image at time *t* + Δ*t*. Fundamentally, this spatial cross-correlation determines ensemble particle displacement between two sequential images (Fig. 2). The farther the particle displacement during the time Δ*t*, the broader the cross-correlation peak will be. In order to quantify the relationship between cross-correlation peak area and the diffusion coefficient, we determine the width of the cross-correlation peak, *s_c_* (pixels) at a height of 1/*e* of the original correlation peak height. We additionally perform autocorrelation on the images. Autocorrelation correlates the interrogation window at time *t* with itself (Fig. 2). The autocorrelation peak is taller and narrower when compared to the cross-correlation peak, with a peak width *s_a_*. Using this information, we calculate the diffusion coefficient (D) using the rearranged equation derived from Olsen and Adrian:^22^

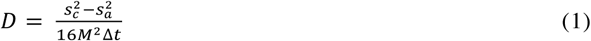

where *M* is the magnification of the microscope objective. Because the peak width has units of pixels, using Equation 1, we can see that the difference in the squared peak widths, 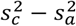, corresponds to the change in the cross-sectional area of the correlation peak at 1/*e*.

**Fig. 2.**
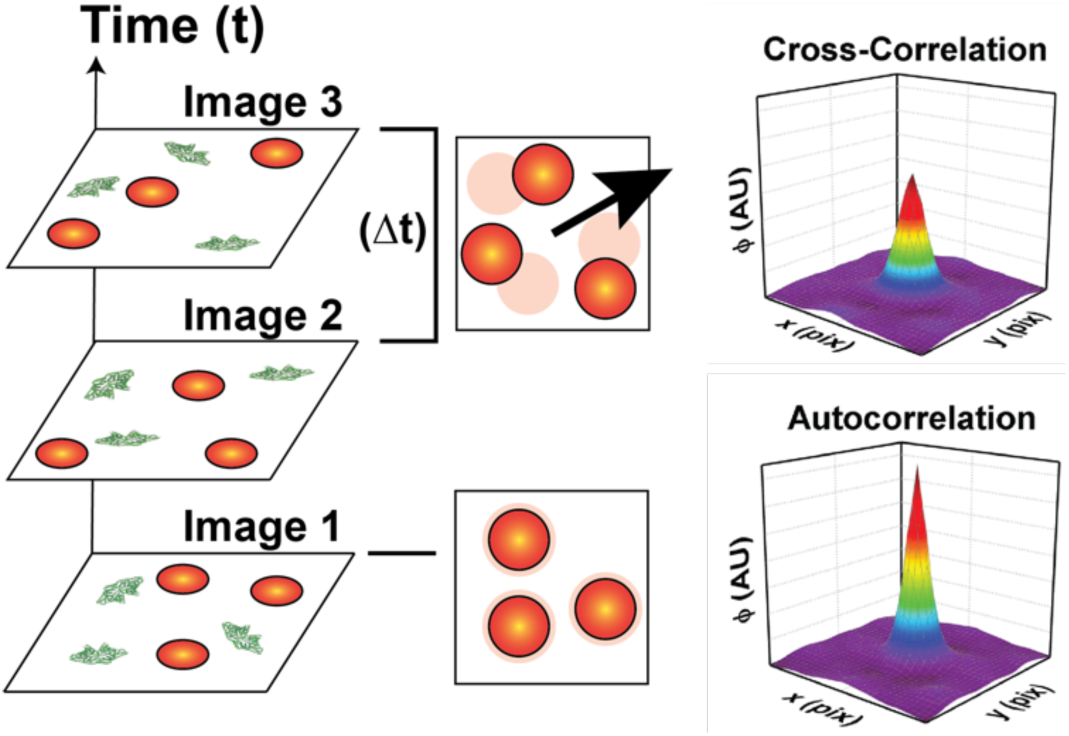
Particle diffusion and image correlation. A stack of images is split into smaller interrogation regions, as pictured. Images which are correlated with themselves produce an autocorrelation peak (Image 1). The correlation of sequential images (Image 2 with Image 3) provide cross-correlation peaks. Note that the cross-correlation peak is both wider and shorter as compared to the autocorrelation peak.

By experimentally determining the diffusion coefficient from the particle images, we can algebraically rearrange the Stokes-Einstein relationship (Equation 2) in order to calculate the viscosity η of a solution according to^23^

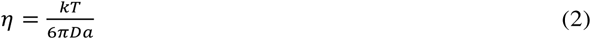

Here, *k* is the Boltzmann constant, *T* is the absolute temperature, and *a* is the hydrodynamic radius of the polystyrene spheres in the protein solution.

We are specifically interested in how the presence of protein and how protein folding state alters solution viscosity. Therefore, we analyze the relative solution viscosities, rather than their actual magnitudes. We determine relative viscosity by algebraic manipulation of Equation 2, where η_0_ is the viscosity of the solution without the solvent protein (but still including the 200 nm polystyrene fluorescent particles).

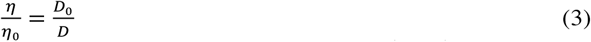

We first measure particle Brownian motion in protein-free solution to determine a baseline viscosity measurement, η_0_. The nanoparticle concentration is specifically chosen to yield statistically relevant results while limiting hydrodynamic interactions. To ensure minimization of any particle-particle interactions that would confound our viscosity measurements, we use the relationship by Batchelor for a dilute monodisperse species of particles,

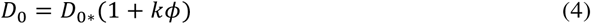

where *D*_0_ is the effective diffusion coefficient from the addition of the polystyrene spheres, *D*_0*\**_ is the diffusion coefficient of a single sphere present in the solvent (where *φ* approaches the 0 limit), *k* is the type-specific constant (we use a value of 2), and *φ* is the volume fraction of the particles in solution.^24–26^ From Equation 4, the percent change in the diffusion coefficient due to the introduction of particles at a concentration of 2.88 x 10^8^ particles/mL is 0.0025%. The hydrodynamic interactions of the particles may be considered negligible when this value is less than 0.01.^26^

PD is fundamentally different than particle tracking, a technique used for studying passive microrheology.^10,27,28^ Particle tracking calculates individual particle trajectories and averages them to determine a diffusion coefficient.^27,29^ Since particle tracking approaches require averaging many particle trajectories to determine statistically relevant results, particle tracking measurements take more time and are computationally more intensive relative to PD.

## Methods

### Preparation of Proteins for Viscosity Measurements

Bovine serum albumin fraction V (BSA, Dot Scientific, Batavia, IL) was suspended in 1X PBS, pH 7.4. All intact samples were maintained at 4°C prior to imaging. To denature BSA, protein samples were heated to 95°C for 2 hours. Additionally, three different insulin solutions were studied in this work. The first insulin solution, from bovine pancreas (Sigma Aldrich, St. Louis, MO, USA), was solubilized in 1X PBS with 1% glacial acetic acid, pH 2.5, in accordance with manufacturer instructions. This insulin sample was used to study protein folding state in acidic conditions. The second insulin sample, also from bovine pancreas, was supplied in 25 mM HEPES, pH 8.2 (Sigma Aldrich, St. Louis, MO, USA), and further diluted in the same buffer. This second insulin solution was used to study insulin state at slightly basic conditions. The third insulin sample was used to study the impact that PBS and HEPES would make on insulin folding state. We lyophilized the insulin (originally stored in 25 mM HEPES) overnight and resuspended the solution in a 1% glacial acetic acid to lower the pH to 2.5. All intact insulin samples were stored at 4°C prior to imaging. Denaturation of insulin occurred by heating samples at 95°C for 2 hours. To determine the sensitivity of the relative viscosity measurements to the amount of insulin degradation in sample, mixtures of intact and denatured insulin at both pH 8.2 and 2.5 were combined volumetrically at 100:0, 90:10, 80:20, 70:30, 60:40, 50:50, 40:60, 30:70, 20:80, 10:90, and 100:0, v/v, denatured:intact at a constant concentration of 6 mg/mL. To study the viscosity of mAb solutions at varying concentrations, 5 mg/ml Rabbit Isotype IgG (Thermo Fisher Scientific, Erie, New York) was snap frozen and lyophilized overnight (FTS Systems Dura-Dry, Stone Ridge, NY). Samples were resuspended in water to a concentration of 50 mg/ml and diluted in 1X PBS to concentrations of 40, 30, 20, and 10 mg/ml.

We used A280 spectral readings to determine all protein concentrations. Each protein sample was measured three times on a Nanodrop 2000 (Thermo Scientific, Erie, NY, USA) and measurements were averaged to determine final protein concentration.

Native polyacrylamide gel electrophoresis (PAGE) was performed as a gold-standard method to determine protein folding state. Protein samples mixed with 4X native loading buffer were introduced to precast polyacrylamide gels (Mini-PROTEAN TGX, Bio-rad, Hercules, CA, USA). Electrophoresis was performed at 120V for 1 hour and 20 minutes in 4°C, followed by staining in GelCode^TM^ blue (ThermoFisher Scientific, Erie, NY, USA) for thirty minutes, and de-stained in deionized water overnight. All protein gels were imaged on a LI-COR Odyssey (Lincoln, NE, USA).

### Particle Diffusometry Viscosity Measurements

200 nm particles (Fluoro-max red dyed aqueous spheres, Thermo Scientific, Erie, NY, USA) were washed in either HEPES pH 8.2 or 1X PBS prior to use by centrifugation at 13,000 x g for 15 minutes. Washed particles were added to protein solutions immediately prior to imaging at a final particle concentration of 2.88 x 10^8^ particles/mL. All protein solutions were stored at 4°C prior to imaging.

A simple fluid well was made by punching a 6 mm diameter through hole (120 μm thickness) in double-sided tape (Therm-O-Web, Wheeling, IL) and adhering the tape to a cover glass slide (Thickness No. 1, Thermo Scientific, Erie, NY, USA). 3 μL of sample (protein solution plus nanoparticles) was introduced to the fluid well and sealed off with a second piece of cover glass, limiting convective evaporation. The sample was imaged using an inverted fluorescence microscope (Nikon TE-2000U, Nikon, Japan) equipped with an X-cite lamp with 40X magnification. Images were recorded using a PCO 1600 CCD camera (PCO, Kelheim, Germany) with an 800 x 800 pixel^2^ imaging window with 2 x 2 binning at 12.5 fps at the vertical middle plane of the chip (to ensure that particle diffusion was unhindered by the glass slides). As this method uses volumetric illumination, all particles in the field of view were imaged, including those in front of and behind the microscope focal plane. However, as particles get farther from focus, their contribution to the correlation function decreases in a known way according to an expression derived by Meinhart *et al*.^30,31^ The effective measurement depth (depth of correlation in PIV literature) is calculated to be 4.2 μm. Particle images were processed and auto and cross-correlation was performed using an in-house MATLAB code in order to determine the diffusion coefficient. 64 x 64 pixel^2^ interrogation windows containing, on average, 8-10 particles were used for 100 image frame stacks (∼8 seconds of data) for a high signal-to-noise ratio while maintaining a statistically relevant number of data points. Nine repetitions, where 100 images constituted an individual measurement, were performed for every individual sample. A two-dimensional Gaussian curve fit was used to calculate the orthogonal profile of the auto- and cross-correlation peaks for both the XZ- and YZ-planes. The width of the correlation peak is defined by 1/e and the width of the XZ- and YZ-Gaussian curves are averaged as one peak width value. To compare all viscosity measurements from the PD measurements, student t-tests were performed between each and every measurement. A 95% confidence interval was used.

### Lower Limit of Detection Measurement

The lower limit of detection (LLOD) was calculated according to the equations found in literature.^32,33^ First, the limit of blank (LOB) was calculated by:

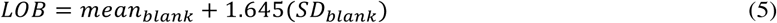

where the *mean*_black_ is the mean value of the viscosity of 200nm particles in buffer (sans protein), and *SD*_black_ is the standard deviation of that same sample. From calculating the LOB we calculate the LLOD as:

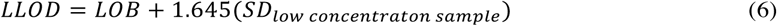

where the *SD_low concentration sample_* is the standard deviation of a low concentration analyte, here being the viscosity measurement of the lowest concentration of the protein measured with PD for every data set. Therefore, our LLOD is expressed as a relative viscosity value.

### Non-Specific Protein Adsorption on Particles

To determine the extent of BSA nonspecific adsorption onto the 200 nm particle surface, particles suspended in all concentrations of the BSA solutions studied (0.01 mg/ml – 10 mg/ml) at a final volume of 100 μL were incubated together for 1 hour. The particle-protein solutions were centrifuged at 13,000 x g and resuspended in 100 μL three times, with a final resuspension in a final volume of 15 μL of 1X PBS. This procedure was performed in triplicate. Particles were then combined with 4X SDS-PAGE loading buffer and boiled at 95°C for 5 minutes. Samples were run on an SDS-PAGE gel for 1 hour and 20 minutes at 120V at room temperature. The SDS-PAGE gel was stained with coomassie (GelCode Blue, ThermoFisher Scientific, Erie, NY, USA) for thirty minutes with gentle rocking followed by destaining in deionized water overnight. Gels were imaged with a LI-COR Odyssey. To compare the levels of protein present in the SDS-PAGE samples, the integrated pixel intensity of each protein band was found using LI-COR Odyssey system software. The integrated pixel intensity was used to back calculate concentrations of protein present in the band by also running known BSA “standards” at concentrations of 0.005, 0.0075, 0.01, 0.05, 0.1, and 0.25 mg/ml. The values for different protein concentrations from the SDS-PAGE gels for denatured and intact protein were compared with a Tukey multiple comparison two-way ANOVA with a confidence level of 95%. The two-way ANOVA investigated how either intact versus denatured protein affected adsorption to the particles as well as how different concentrations affect protein adsorption.

### FITC Staining of Protein on Particles

Protein-particle solutions were stained in Fluorescein Isothiocyanate (FITC) to visually confirm protein content in the particle solution under fluorescence microscopy. FITC was dissolved at 1 mg/ml in DMSO prior to staining. Particle-protein solutions were adjusted to 0.1M sodium carbonate. The 1 mg/ml dissolved FITC was added to the protein-particle solution at a 1:20 v/v, respectively, and incubated in the dark by rotation for 8 hours at 4°C. Following, NH_4_Cl was added to the aliquots to a final volume of 50mM and incubated in the dark by rotation for 2 hours at 4°C. For washing off excess protein to analyze nonspecifically adsorbed protein on the particle surface, particles were washed three times by centrifugation at 13,000 x g and suspended to their same initial volume in 1X PBS. The fluorescent stained samples were imaged using the inverted fluorescence microscope (Nikon TE-2000U, Nikon, Japan) equipped with an X-cite lamp with 40X magnification on an Alexa 488 filter cube. Images were recorded with a DS2 camera (Nikon, Japan) and NIS Elements software (Nikon, Japan).

## Results and Discussion

### Investigating the Viscosity of BSA Solutions

BSA is used as a model protein to perform protein viscosity characterization studies in the PD system. The two initial parameters of interest are the effect of (1) concentration and (2) folding state of BSA on solution viscosity. To investigate the differences between intact and denatured BSA, we perform a native PAGE on solutions of BSA at 0.25 mg/mL with and without heat treatment (Fig. 3A). As a consequence of crosslinking that occurs among semiflexible polypeptide chains during heat denaturation, the denatured BSA is not electrophoretically mobile as it is too large to penetrate through the polyacrylamide gel.^34^ In contrast, intact BSA displays several distinct bands at molecular weights that likely correspond to the presence of monomers, dimers, and oligomers in the BSA solution.^35^ A solution of denatured and intact BSA solution at a 1:1 v/v ratio shows features of both electrophoretically immobile denatured BSA and the presence of BSA monomers, dimers, and oligomers (Fig. 3A).

**Fig. 3.**
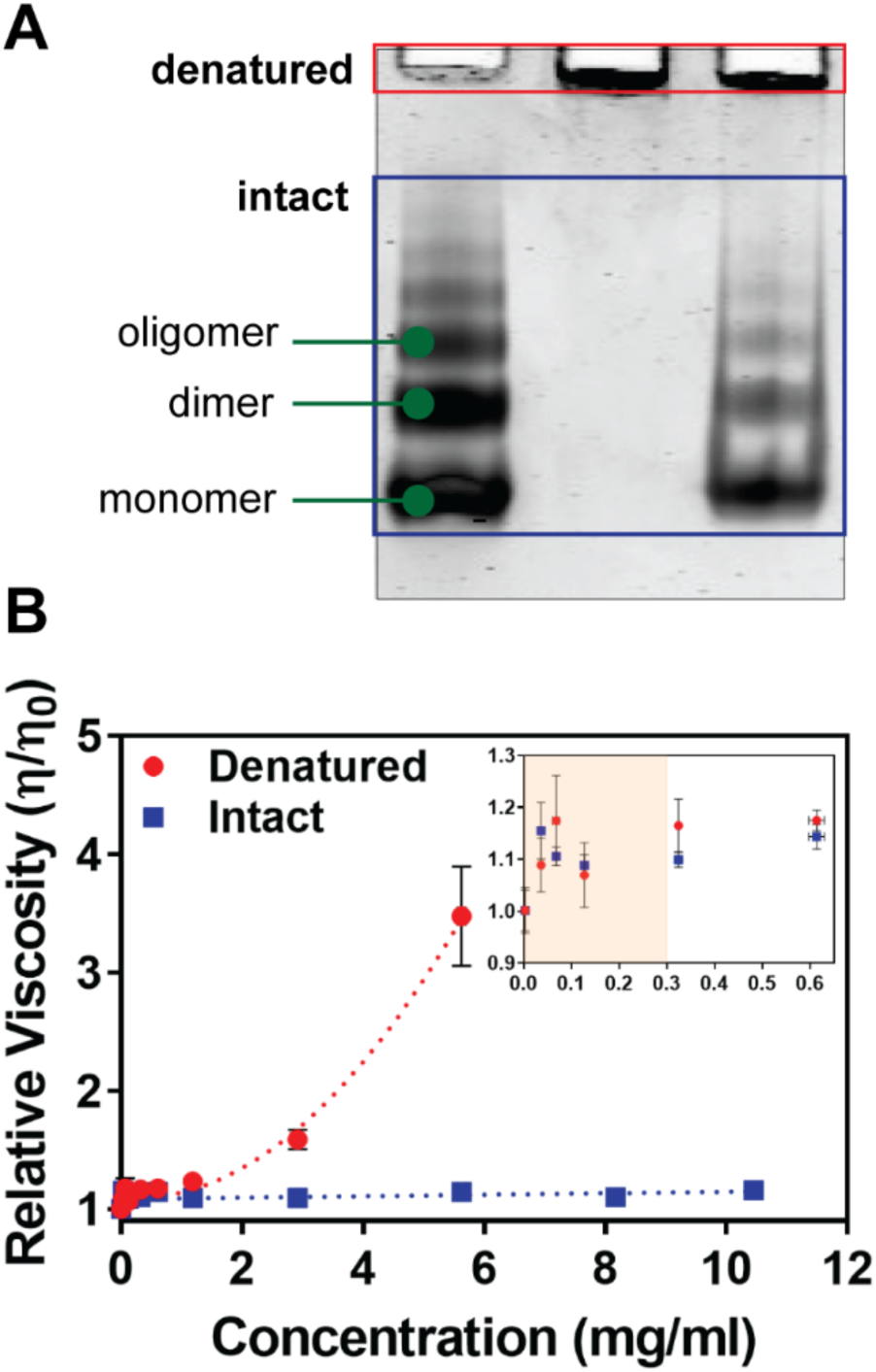
Intact and denatured BSA solutions. (A) Native PAGE of intact and denatured BSA at a concentration of 0.25 mg/ml. The denatured BSA remains at the channel entrance of the gel (surrounded by a red box) and intact samples propagate into the gel (blue box). The left lane contains only intact BSA, the middle lane contains only heat treated, denatured BSA, and the third, rightmost lane contains a mixture of 50% native and 50% degraded BSA. (B) PD was used to measure the relative viscosity of BSA solutions. The relative viscosity of denatured BSA solutions (red circles) increases as the concentration of the protein increases, whereas the viscosity of solutions containing intact protein (blue squares) remains constant as a function of concentration. The viscous effects from lower protein concentrations are statistically indistinguishable from one another as both a function of concentration and protein folding state (inset, highlighted in the peach region). n = 9, see Supplemental Information for raw data.

Measurements of the relative viscosity of solutions of BSA with and without heat treatment were performed using PD (Fig. 3B). Measurements of heat denatured BSA solutions could only be performed up to a concentration of 5 mg/ml as gelling occurred in samples above this concentration, causing significant errors in pipetting. We observe that the viscosity of denatured BSA solutions dramatically increases as a function of concentration (Fig. 3B). PD can be used to determine differences in viscosity between solutions of intact and denatured BSA at concentrations of approximately 0.3 mg/ml and greater (p < 0.001 for 0.3 and 0.6 mg/ml and p < 0.0001 for 1 mg/ml and greater), raw PD data in Tables S1† and S2†).

For intact BSA, we observe no increase in the solution viscosity at increasing BSA concentrations (p-value > 0.05). We calculate a lower limit of detection (LLOD) of BSA viscosity measurements with PD to be at a relative viscosity of 1.12, in this case meaning that PD can measure the viscosity of denatured BSA at concentrations of 0.3 mg/ml and greater. Individual native BSA proteins, like other globular proteins, can be modeled as hard rigid spheres moving in space. On the other hand, denatured BSA, similarly to other denatured proteins, is likely better-described as a discrete semiflexible polymer.^36–38^ Thus we speculate that the non-linear trend of increasing viscosity with increasing concentration of denatured protein in Fig. 3B is likely due to increases in unfolded protein crosslinking at increasing concentrations.

### Non-Specific Protein Adsorption on Viscosity Measurements

The relative change in solution viscosity that we calculate in Fig. 3B may not be a function of protein denaturation alone. Particles without chemical surface modifications are likely to have non-specific adsorption of proteins to their surfaces, thus increasing the particles’ hydrodynamic radii.^17^ As a particle’s radius increases, its diffusion coefficient decreases according to the Stokes-Einstein equation. Therefore, the effect of low levels of protein adsorption onto particle surfaces, even at low protein concentrations and regardless of protein folding state, may contribute to lower sensitivity of the PD measurement.

To study the effect that non-specific adsorption of proteins onto our particles has on PD measurements, we first investigate whether or not intact BSA non-specifically adsorbs onto unmodified particle surfaces. Polystyrene particles were incubated with fluorescently labeled FITC-BSA, imaged, washed to remove the excess BSA, and imaged again to visualize any remaining fluorescent protein that is attached to the particle surface (Fig. 4A). In Fig. 4A it is evident that in the unwashed sample FITC-BSA is dispersed throughout, as indicated by green fluorescence (FITC). After removing the free BSA, the FITC signal is localized to the particle surfaces. This confirms that non-specific adsorption of BSA is, in fact, occurring on these unmodified red fluorescent polystyrene spheres.

**Fig. 4.**
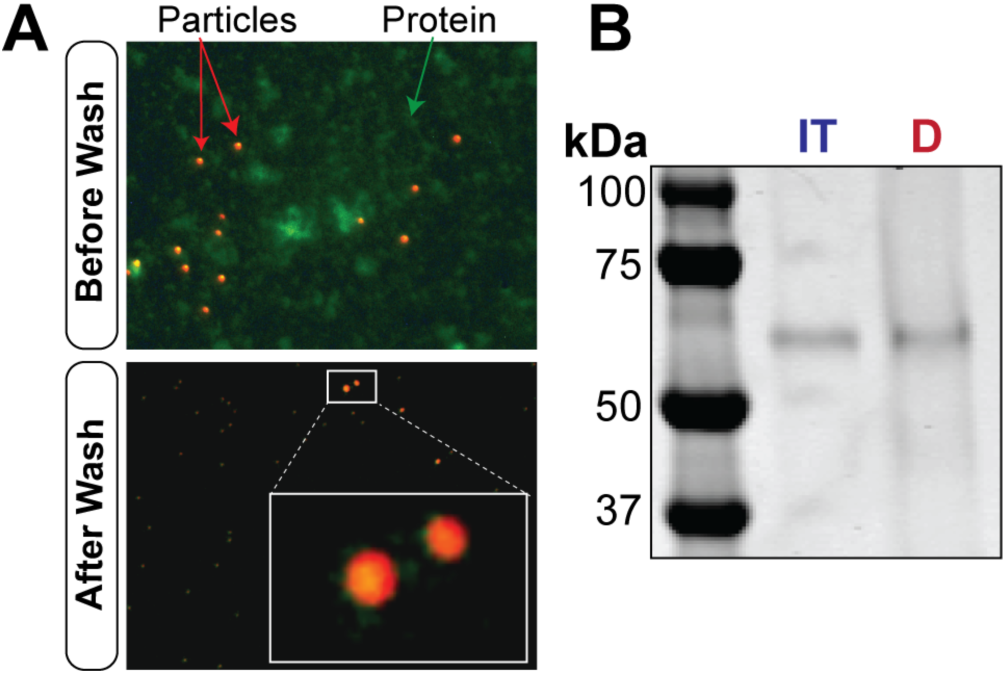
Nonspecific protein adsorption on particle surfaces. (A) Prior to washing, particles in the presence of 5 mg/ml FITC labeled BSA show green fluorescent background signal, indicating free protein (top). After washing (bottom) the background fluorescent signal is dramatically reduced as expected, with concentrated fluorescent green signal located around the red particle circumference indicating non-specific adsorption of FITC-BSA to the particles. (B) A representative SDS-PAGE analysis of the 5 mg/ml BSA sample non-specifically absorbed to beads for both intact and denatured BSA samples indicate similar levels of non-specific adsorption.

We perform a semi-quantitative SDS-PAGE analysis to determine how much BSA non-specifically adsorbed to the particle surface. Particles are incubated with either intact or denatured BSA solutions at varying protein concentrations (0.01 – 10 mg/ml), washed, boiled in the presence of SDS, and analyzed. SDS-PAGE analysis of the boiled protein-particle samples shows that the protein can be stripped from the particles and visualized with coomassie (Fig. 4B). From the SDS-PAGE, the integrated pixel intensities of all the protein bands are measured with image processing (gel images are in the Fig. S1†, values in Table S3†) and are compared using a two-way ANOVA with a post-hoc Tukey test to determine if protein folding state or protein concentration have an effect on non-specific adsorption onto the particle surface. We find no statistically significant difference (p-value > 0.05) among the integrated pixel intensity values of the protein bands for intact BSA and denatured BSA, respectively at all concentrations. This indicates that protein folding state does not change the amount of protein non-specifically adsorbed to the particle. Furthermore, the integrated intensity values of the SDS-PAGE bands at all concentrations (0.01-10 mg/ml) relative to one another are also not statistically significantly different (p-value > 0.05), indicating that concentration does not play a role of the quantity of non-specific protein adsorption onto the particles. Together we take these results to mean that similar amounts of protein non-specifically adsorb onto particle surfaces regardless of folding state (denatured vs intact protein) or concentration of protein in solution. Thus, we assume that all particles, regardless of protein treatment, undergo the same surface adsorption, and the differences in diffusion coefficient that we measure with PD indicates changes in the viscosity of the protein-particle solutions, rather than differences in particle size due to non-specific adsorption. This also indicates that differences in solution viscosity at lower protein concentrations maybe obtainable with particle surface modifications that are meant to block non-specific adsorption.

### Viscosity Measurements of Insulin Solutions

Having determined that PD can readily determine differences of viscosity between intact and denatured BSA solutions, we sought to characterize the change in solution viscosity of intact and denatured state of a more pharmacologically relevant protein, insulin. Insulin has vastly different physiochemical properties than BSA. The molecular weight of insulin is 5.8 kDa (BSA is 66.5 kDa, SDS-PAGE gel in Fig. S2†) and is a heterodimer comprised of separate *α* and *β* subunits.^39^ When insulin is denatured, the protein separates into its two respective subunits due to the breakage of disulfide bonding separating into two 21 and 30 amino acid polypeptide chains, respectively.^40^ In contrast, BSA when unfolded is a single 607 amino acid polypeptide chain.^41^

There are currently multiple formulations of insulin produced by biopharmaceutical companies. As many of these formulations are proprietary and the exact formulation is not public information, we measured the viscosity of insulin solutions comprised of different buffers (HEPES and PBS) at two different pH values (2.5 and 8.2). Proper folding of insulin in PBS pH 2.5 was verified by native PAGE (see Fig. S3†). The relative viscosities of intact and degraded insulin at varying concentrations are measured using PD (Fig. 5A-C). We observe that, similar to BSA, the viscosity of denatured insulin solutions dramatically increases as the concentration of the protein increases (raw data in Table S4†). From this the LLOD of insulin in PBS is at a relative viscosity of 1.09, which is similar to our measurement for BSA LLOD (1.12). This relative viscosity measurement occurs at a concentration of denatured insulin in PBS pH 2.5 somewhere below 2 mg/ml, indicating that we can detect significant differences in the viscosity of denatured compared to intact insulin at concentrations of 2 mg/ml and greater (p < 0.0001, Fig. 5A), but not below.

**Fig. 5.**
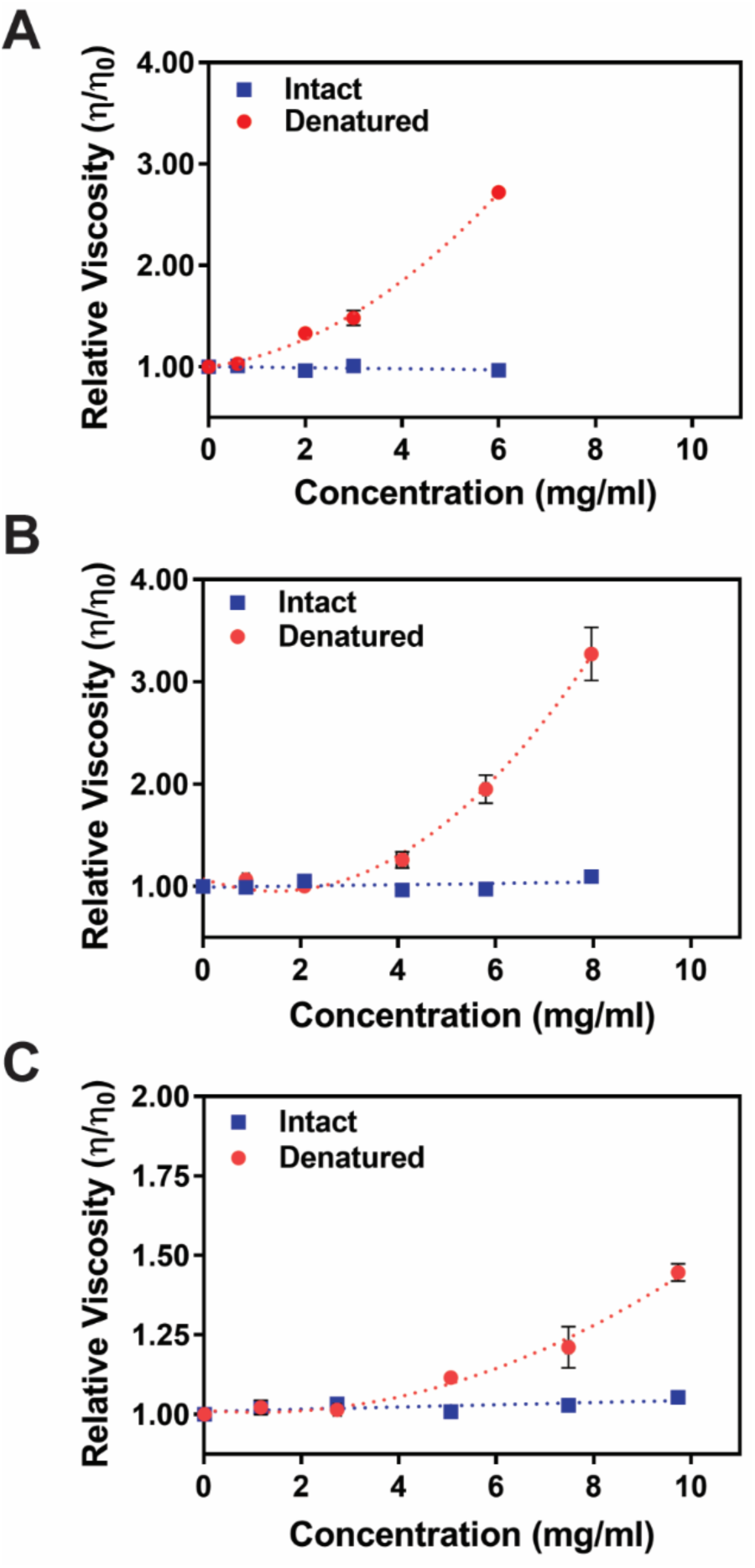
The relative viscosity of insulin solutions as a function of concentration. (A) PBS pH 2.5. There is a dramatic difference in the viscosity of denatured insulin at 2 mg/ml as compared to intact insulin whereas in (B) the viscosity of denatured insulin solubilized in HEPES pH 2.5 shows a statistically significant difference from the intact samples at a concentration of 4 mg/ml. (C) Insulin samples in HEPES pH 8.2 shows the least dramatic difference between intact and denatured insulin solutions at a concentration of 5 mg/ml but still shows a similar trend from (A) and (B) in that there is a non-linear increase in solution viscosity for denatured protein sample. Note that the y-axes in (A) and (B) are different than (C). n=9.

Interestingly, changes in the relative viscosity of denatured insulin as a function of concentration varies with buffer and pH. After performing the insulin viscosity study in PBS, PD is used to assess the viscosity of insulin in HEPES pH 2.5. Significant differences in the viscosity of denatured and intact insulin are detectable at concentrations of 4 mg/ml and greater when solubilized in HEPES at pH 2.5 (Fig. 5B). The LLOD of PD measurements of the relative insulin viscosity of HEPES pH 2.5 is 1.07, which occurs at the 4 mg/ml denatured protein concentration. PD measurements of relative viscosity of intact insulin in HEPES pH 2.5 do not significantly vary as a function of concentrations (PD data in Table S5†). Together with the LLOD measurements of BSA and insulin in PBS pH 2.5, these results suggest that PD can be used to measure changes in relative viscosity at least greater than 0.07 of any sample (BSA, insulin, and other biomolecules).

We also measured the relative viscosity of insulin solutions in HEPES pH 8.2. Measured changes in relative viscosity of denatured insulin show a much less drastic non-linear increase in viscosity at increasing concentrations as compared to the PBS and HEPES pH 2.5 cases. In the HEPES pH 8.2 buffer condition, PD measures the difference in viscosity between intact and degraded insulin at a concentration of 5 mg/mL and greater (p < 0.0001, Fig. 5C) which is where PD reaches its LLOD. Additionally, following the same trend as the PBS and HEPES pH 2.5 cases, there is no discernible change in relative viscosity between all concentrations of intact insulin in HEPES pH 8.2 over the range of concentrations measured (PD data in Table S6†). Taking these results together, we see that that denatured insulin solubilized in buffers which are closer to physiological pH have less dramatic measurable changes in viscosity. Likewise, the buffer/salt content, such as PBS versus HEPES, has an effect on the viscosity of denatured insulin solutions as well, suggesting that PD can detect how the solubility of denatured protein changes as a function of changes in buffer conditions.

We attribute these differences in changes in relative viscosity to the interaction of the denatured protein with the various ions and salts in the different buffer and pH solutions. Ion and salt content of solutions are known to affect protein morphology, interacting with the exposed amino acid side chains of the proteins to either shield or change individual amino acid charge. The isoelectric point of insulin is pH 5.3; thus the overall charge of insulin is positive in the acidic pH and negative in the basic buffer. These molecular interactions are likely having a direct effect on how the denatured polypeptide chains of insulin are interacting with each other in solution, and in turn, affecting measurements of solution viscosity obtained with PD. We conclude that buffer conditions play a major role in the viscosity of denatured protein solutions, and speculate that different biopharmaceutical formulations of insulin would have different absolute viscosity measurements. Regardless, as the concentration of degraded insulin in a sample increases, the relative viscosity of the solution is expected to markedly increase to detectable levels.

### Mixtures of Insulin

Prescription insulin is stored at concentrations between 100 units/ml^42^ to 200 units/ml,^43^ which is equivalent to 3.5-7 mg/ml, assuming all insulin is intact at the time of manufacturing and packaging. However, it is unlikely in practice that prescription insulin will be either 100% degraded or 100% intact. We therefore sought to measure the changes in relative viscosity of mixtures of varying ratios (v/v) of intact and denatured insulin at a consistent concentration of 6 mg/ml (to remain within the range of prescription insulin). Like BSA (Fig. 3A) most denatured insulin does not enter the PAGE gel, and remains at the entrance to the gel channel (Fig. 6A, red box), indicating cross-linking of denatured protein. In contrast, the intact protein is electrophoretically mobile and moves into the PAGE gel (Fig. 6A, blue box). Consistent with the varying ratio of intact to denature insulin, higher intensity bands within the gel channel are present in insulin mixtures containing larger ratios of native insulin, whereas higher intensity bands at the channel entrance correlate with larger volumes of denatured insulin.

**Fig. 6.**
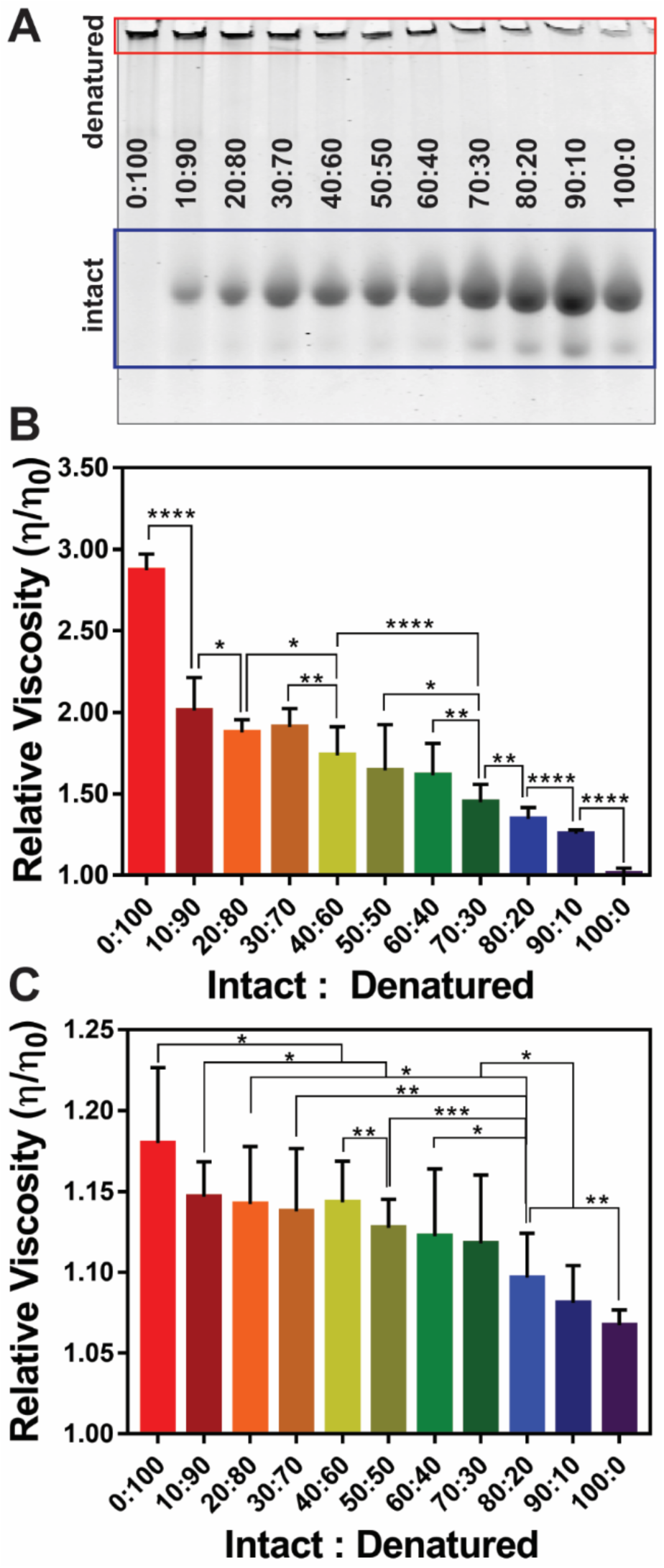
Relative viscosity of insulin mixtures. (A) A native PAGE of different mixtures of intact and denatured insulin. (B-C) The relative viscosity of mixtures of intact and denatured insulin (v/v ratios). (B) Insulin at PBS pH 2.5 and (C) insulin in HEPES pH 8.2. As the ratio of denatured insulin increases, the relative viscosity also increases. Note the relative viscosity on the y-axes is different between (B) and (C) (* indicates p < 0.05, ** p < 0.01, *** p < 0.001, **** p <0.0001, n = 9).

To determine the relative viscosity of the varying insulin mixtures PD measurements of insulin mixtures is performed in both PBS pH 2.5 and HEPES pH 8.2. As expected, the relative viscosity of insulin solutions increases as the percentage of denatured insulin increases (Fig. 6B-C). Insulin solutions in PBS pH 2.5 (Fig. 6B) show a more dramatic change in protein viscosity between each v/v ratio when compared to insulin in HEPES pH 8.2 (Fig. 6C). This behavior is expected given our findings comparing exclusively denatured to exclusively intact insulin solutions (Fig. 5B and 5C). There is a significant difference in the relative viscosity of PBS insulin solutions starting at as little as 10% denatured insulin (p-value = 1E-19). From this, we calculated our lower limit of detection of insulin in PBS to be below the 90:10 intact to denatured insulin mixture. The relative viscosity change is 0.21, compared to the 100:0 intact to denatured insulin mixture (PD data found in Table S7†). In solutions of insulin in HEPES at pH 8.2 we detect a difference in relative viscosity of solutions with 80:20 intact to denatured insulin mixture (relative viscosity change of 0.1, PD data found in Table S8†).

### Monoclonal Antibody Viscosity

Monoclonal antibodies are stored at high concentrations for patient injection. With 70 mAb drugs expected to be on the market by 2020,^9^ there is an opportunity to design high throughput viscosity measurement systems for formulation development and upscale manufacturing. Antibodies have molecular weights typically around 150kDa (with 2 light chains and 2 heavy chains)^44^, which is much larger than BSA and insulin. Because of this, intact antibodies are more likely to cause changes in concentration than smaller biotherapeutic proteins. Therefore, we use PD to study the effect of increasing concentration of intact IgG antibodies on solution viscosity (raw data presented in Table S9†). As expected, there is a gradual increase in solution viscosity as the concentration of antibodies is increased (Fig. 7). Performing a t-test between each concentration, there is statistically significant differences in the relative viscosity of each concentration measured (p < 0.0001 and p < 0.001). A change in concentration of around 10 mg/ml produces changes in solution viscosity of around 0.14 ± 0.02, which is above our LLOD. This is promising because mAb therapeutics remain far above this concentration. Therefore, PD can be used to measure high concentrations of antibody solutions for downstream applications in mAb formulation characterization.

**Fig. 7.**
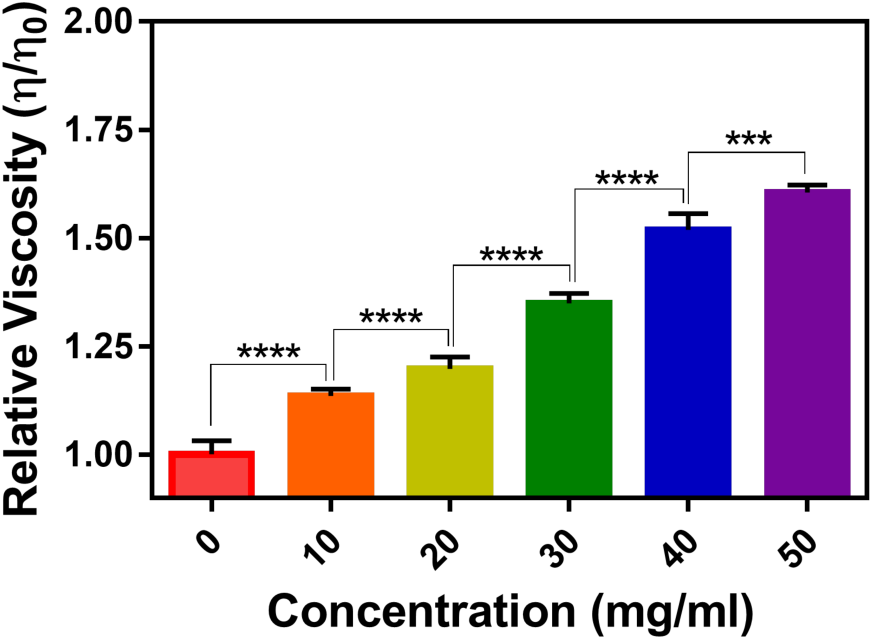
Antibody viscosity. As the concentration of IgG antibody solution increases, the relative viscosity of the solution increases (**** p < 0.0001, *** p < 0.001, n = 9).

## Conclusions

Using particle diffusometry (PD), we have developed a method with which we can determine the degree of protein degradation. With PD we observe that intact proteins show little-to-no change in viscosity up to concentration changes of approximately 10 mg/ml (Fig. 3, 5 and 7). This limit is likely to vary most with the size of the protein. However, heat denaturation of protein solutions produces measurable changes in sample viscosity as a function of increasing concentration (Fig. 3 and 5). This change in the viscosity occurs due to unfolding, crosslinking and aggregation of proteins during the denaturation process. As such, we observe that the concentration at which denatured insulin solutions exhibit viscosities significantly different from solutions with intact protein is a function of solution buffer and pH (Fig. 5). One important implication of these results is that quantitative measurements of protein degradation would require standards and controls with similar buffer formulations to be accurate. Regardless, these measurements are robust and allow for detection of as little as 10% denatured insulin in some formulations (Fig. 6). These results are readily translated to other important biotherapeutic products, such as monoclonal antibodies, a significantly larger protein than insulin (Fig. 7) that are regularly administered to patients at high concentrations. As such, the viscosity of mAb solutions is a critically important parameter that affects the dosing strategy. Our measurements use sample volumes of 3μL and imaging times of 8 seconds. Our current algorithms implemented on an Intel^®^ Core^™^i5-3230M CPU at 2.60GHz computer processer require approximately 18 seconds per sample. Thus time-to-result per sample is as low as 30 seconds and could be optimized to run even faster. A tool like PD would enable rapid, high throughput, and low volume measurements of biotherapeutic formulations, and may be implemented for formulations research and development, or in manufacturing and distribution settings. We also envision PD-enabled point of care diagnostics for clinics and patients. Integration of PD with small handheld devices would enable pharmacists and patients to track the viability of their protein-based prescriptions. Implementation of particle diffusometry for measuring the viscosity of biotherapeutic solutions could be used to optimize pharmaceutical formulations, track biotherapeutic stability throughout the manufacturing and distribution chain, and be used in clinical settings as measure of the efficacy of a biotherapeutic. Implementing these methods could potentially decrease prescription waste, decrease incorrect drug use, mitigate adverse reactions in patients, and provide new means for patients to control their own health and prescription monitoring.

## Acknowledgements

We acknowledge support from the Purdue Office of the Executive Vice President for Research and Partnerships.

